# A H3K9me2-Binding Protein AGDP3 Limits DNA Methylation and Transcriptional Gene Silencing in Arabidopsis

**DOI:** 10.1101/2021.03.08.434320

**Authors:** Xuelin Zhou, Mengwei Wei, Wenfeng Nie, Yue Xi, Xuan Du, Li Peng, Qijie Zheng, Kai Tang, Viswanathan Satheesh, Yuhua Wang, Jinyan Luo, Rui Liu, Zhenlin Yang, Yingli Zhong, Guo-Yong An, Jian-Kang Zhu, Jiamu Du, Mingguang Lei

**Affiliations:** Shanghai Center for Plant Stress Biology, CAS Center for Excellence in Molecular Plant Sciences, Chinese Academy of Sciences, Shanghai 201602, China; Key Laboratory of Molecular Design for Plant Cell Factory of Guangdong Higher Education Institutes, Institute of Plant and Food Science, School of Life Sciences, Southern University of Science and Technology, Shenzhen 518055, China; University of Chinese Academy of Sciences, Beijing 100049, China; Department of Horticulture & Landscape Architecture, Purdue University, West Lafayette, IN 47906, USA; State Key Laboratory of Crop Stress Adaptation and Improvement, School of Life Sciences, Henan University, Kaifeng 475004, China

**Keywords:** AGDP3, ROS1, DNA demethylation, H3K9me2, Epigenetics

## Abstract

DNA methylation is critical for tuning gene expression to prevent potentially deleterious gene-silencing. The Arabidopsis DNA glycosylase/lyase REPRESSOR OF SILENCING 1 (ROS1) initiates active DNA demethylation and is required for the prevention of DNA hypermethylation at thousands of genomic loci. However, the mechanism recruiting ROS1 to specific loci is not well understood. Here, we report the discovery of Arabidopsis AGENET Domain Containing Protein 3 (AGDP3) as a cellular factor required for ROS1-mediated DNA demethylation, and targets ROS1 to specific loci. We found that AGDP3 could bind to the H3K9me2 mark by its AGD12 cassette. The crystal structure of the AGDP3 AGD12 in complex with an H3K9me2 peptide reveals the molecular basis for the specific recognition, that the dimethylated H3K9 and unmodified H3K4 are specifically anchored into two different surface pockets. Interestingly, a histidine residue located in the methylysine binding aromatic cage enables AGDP3 pH-dependent H3K9me2 binding capacity. Considering the intracellular pH correlates with the histone acetylation status, our results provide the molecular mechanism for the regulation of ROS1 DNA demethylase by the gene silencing H3K9me2 mark and the potential crosstalk with active histone acetylation mark.

## Introduction

DNA methylation, characterized by adding a methyl group onto the fifth position of the cytosine, has profound impact on biological processes such as gene regulation, transposable element silencing and genome stability (Robertson, 2005; Slotkin and Martienssen, 2007; Zhang et al., 2018b). In plants, DNA is methylated at specific loci by DNA methyltransferases controlled by the RNA-directed DNA methylation (RdDM) pathway (Law and and Jacobsen 2010; Matzke and Mosher, 2014). Once established, different mechanisms are required to maintain this epigenetic mark, depending on the cytosine context. While the methylation in the symmetric CG and CHG (where H is C, A or T) context is copied during DNA replication by DNA methyltransferase 1 (MET1) and chromomethylase CMT3, respectively, the asymmetrical CHH methylation is *de novo* methylated by domain rearranged methyltransferase 2 (DRM2) through RdDM pathway and by CMT2 (Lindroth et al., 2001; Matzke et al. 2009; Zemach et al., 2013). On the other hand, DNA methylation can be actively erased by a class of bifunctional DNA glycosylases/lyases, including REPRESSOR OF SILENCING 1 (ROS1), DEMETER (DME), DME-LIKE 2 (DML2), and DML3 (Zhu, 2009). While DME is preferentially expressed in companion cells of gametes and functions in genomic imprinting (Gehring et al., 2006; Hsieh et al., 2009; Huh et al., 2008; Penterman et al., 2007), DML2, DML3, and ROS1 mainly function in locus-specific DNA demethylation and in preventing transcriptional silencing of endogenous and transgenic loci (Gong et al., 2002; Penterman et al., 2007; Qian et al., 2012).

ROS1 is targeted to specific genomic loci through the Increased DNA Methylation (IDM) complex comprising IDM1, IDM2, IDM3, methyl CpG-binding protein 7 (MBD7) and Harbinger transposon-derived proteins HDP1 and HDP2 (Qian et al., 2012; Qian et al., 2014; Lang et al., 2015; Duan et al., 2017). MBD7 preferentially binds to densely methylated CG regions and recruits IDM1 to acetylate histone H3K18 and H3K23 to facilitates ROS1-mediated DNA demethylation (Qian et al., 2012; Lang et al., 2015). At a subset of H2A.Z-enriched loci, MBD9 and SNX1 recognize these acetylated histone and recruit SWR1 complex to deposit H2A.Z, which tethers ROS1 to demethylate DNA methylation (Nie et al., 2019; Sijacic et al., 2019). Recently, it was reported that another methyl DNA binding protein RMB1 can interact with ROS1 to regulate DNA methylation at several loci, independently of the IDM protein complex (Liu et al., 2020).

DNA is compacted around the histone octamer, which can also be covalently modified at the N-terminal tails. These post-translational modifications can change chromatin states and/or recruit some histone readers, which may further modify the histone and even the DNA. For example, H3K9me2, a mark rich in pericentromeric heterochromatin regions, is associated with transcriptional repression. It is created by H3K9 methyltransferases KRYPTONITE, SUVH5 and SUVH6, which are recruited mainly by binding CHG methylation. On the other hand, CMT2 and CMT3 can recognize H3K9me2 and are recruited to methylate CHH and CHG, respectively. Thus, H3K9 methylation and DNA methylation form a feedback loop to reinforce the repression (Du et al., 2015). AGENET domain (AGD) belongs to the ‘Royal Family’ histone readers (Maurer-Stroh et al., 2003). In human, the tandem AGDs of FMRP is able to specifically recognize the H3K79me2 mark (Alpatov et al., 2014; Ramos et al., 2006). In plants, the tandem AGDs of AGENET DOMAIN CONTAINING PROTEIN 1 (AGDP1, also known as ADCP1) can specifically recognize the H3K9me2 mark *in vitro* and *in vivo* (Zhang et al., 2018a; Zhao et al., 2019).

AGDP1 is concentrated in centromeric and pericentromeric regions and recruits some unknown factors to regulate non-CG methylation and transcriptional gene silencing (Zhang et al., 2018; Zhao et al., 2019). However, H3K9me2 methylation is also present in facultative heterochromatin, where the expression of resident genes or nearby genes are dynamically regulated through the life cycle or in response to environmental stimuli. We, therefore, opine that there exists some factor(s) that recognize this repressive mark and then change the modification or recruit some other regulator(s) to prevent transcriptional silencing.

Here, we identified an anti-silencing factor AGENET DOMAIN CONTAINING PROTEIN 3 (AGDP3) with two pairs of tandem AGDs. We found that AGDP3 prevents transgenes and some endogenous loci from hypermethylation. We further found that AGDP3 can specifically recognize the gene silencing H3K9me2 mark by its first tandem AGD cassette. Our structural studies revealed the molecular mechanism of the specific recognition of H3K9me2 by AGD12 of AGDP3 and further identified AGDP3 as a pH-dependent histone reader with lower binding in low pH. The intracellular pH is correlated with the histone acetylation status. Therefore, our studies revealed the molecular basis for the specific targeting of ROS1 anti-silencing machinery to its substrate methylated DNA loci, which is enriched with H3K9me2 and further shed light on the mechanism underlying the crosstalk among DNA methylation, H3K9me2 and histone acetylation.

## Results

### AGDP3 prevents gene silencing

To identify potential anti-silencing factors in Arabidopsis, we performed a forward preestablished genetic screen with transgenic plants expressing the cauliflower mosaic virus 35S promoter-driven sucrose transporter 2 *(35S::SUC2).* Overexpression of *SUC2* results in sucrose over-accumulation and seedlings have short roots when grown on medium containing 1% sucrose compared with Col-0 wild-type plants (Lei et al., 2014). We isolated two mutant alleles on sucrose-containing media, *agdp3-1* and *agdp3-2,* displaying long-root phenotype compared with *35S::SUC2* (hereafter referred to as WT) plants (Fig. 1a). Consistent with the root phenotype, the *SUC2* gene expression was silenced in *agdp3-1* mutant (Fig. 1b). The WT also expressed hygromycin phosphotransferase II *(HPTII)* gene driven by double *35S (2×35S)* promoter and conferred hygromycin resistance (Lei et al., 2014). This transgene was also silenced in *agdp3-1* and *agdp3-2* mutants, showing hygromycin sensitive phenotype (Fig. 1a and 1c). Then, we compared the transcriptomes, and found that 51 endogenous genes were significantly downregulated in *agdp3-1* mutant when compared with the WT (FDR=0.05, FC>2) (Fig. 1d). GO enrichment analysis indicated that AGDP3 mutation may affect some biological processes (Fig. 1d). These results show that AGDP3 is required for preventing silencing of both transgenes and some endogenous genes.

**Figure. 1.**
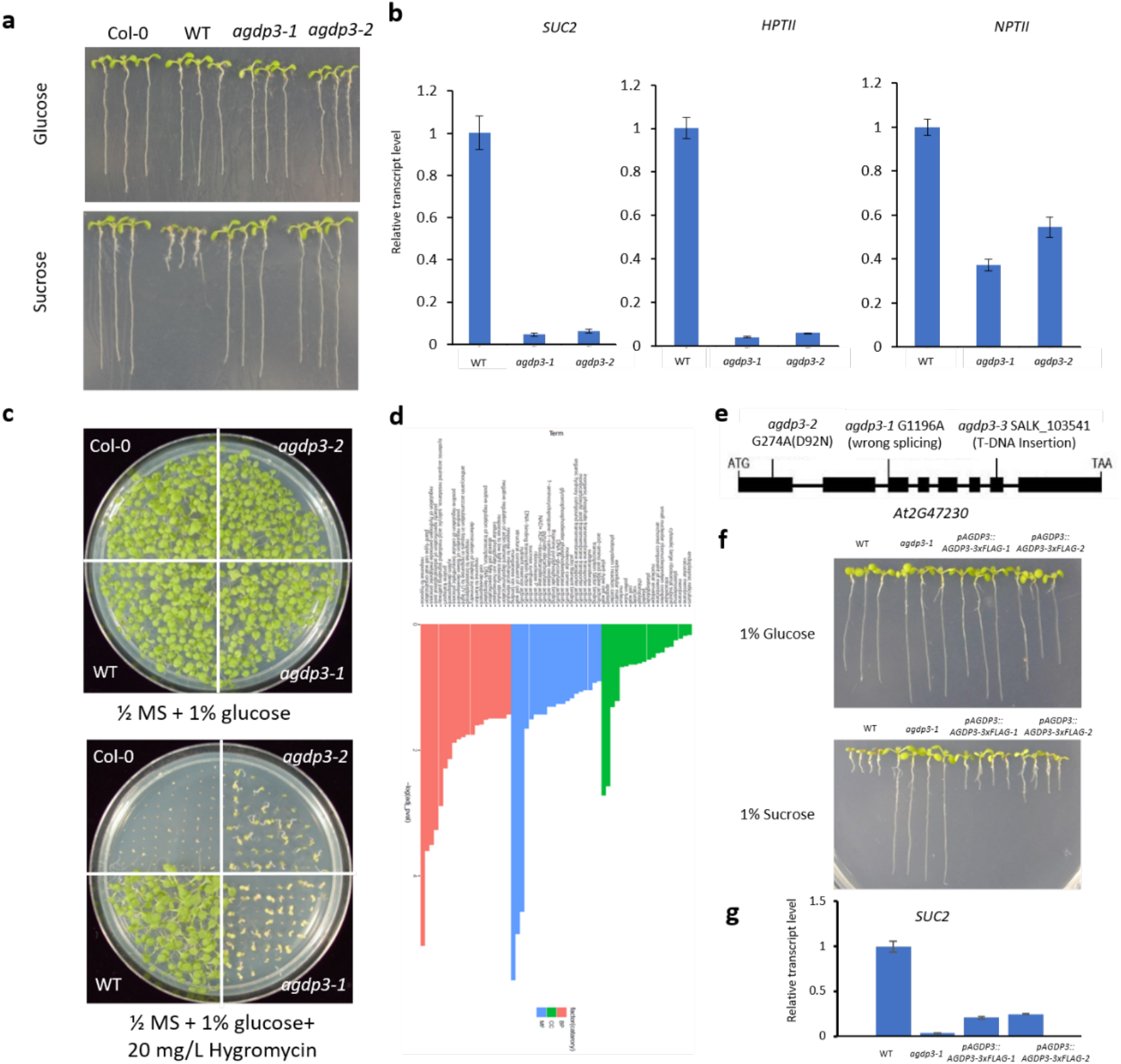
AGDP3 Prevents DNA Hypermethylation and Transcriptional Gene Silencing. **a** Comparison of root growth phenotype among Col-0, 35S::SUC transgenic line (WT), and *agdp3* mutants grown on glucose or sucrose medium. **b** Transcript levels of *SUC2, HPTII* and *NPTII* transgenes in *agdp3-1* and *agdp3-2* mutants compared to WT detected by RT-qPCR. **c** AGDP3 dysfunction causes compromised hygromycin resistance in the transgenic plants. **d** RNA-Seq analysis of WT and *agdp3-1* seedlings. **e** A diagram of the *AGDP3* gene showing the mutation in *agdp3-1* and *agdp3-2* mutant, and the T-DNA insertion position of *agdp3-3* allele. Boxes and lines represent exons and introns, respectively. **f** Expression of *AGDP3* driven by its native promotor rescued the short root phenotype in *agdp3-1.* **g** *SUC2* transcript level in WT, *agdp3-1* and both complementation transgenic lines detected by RT-qPCR.

### AGDP3 regulates DNA demethylation

Through map-based cloning, we found that *agdp3-1* has G to A mutation at the end of the second intron of At2G47230 causing a splicing defect. Also, a G to A mutation in the first exon changes Asp92 to Asn in *agdp3-2* (Fig. 1e). To confirm that the *agdp3* mutations were responsible for the silencing of *35S::SUC2,* the expression of *AGDP3* fused with a *3xFLAG* tag driven by its native promoter *(pAGDP3::AGDP3-3xFLAG)* in *agdp3-1* restored *35S::SUC2* gene expression and rescued the short root phenotype (Fig. 1f and 1g).

AGDP3 was predicted to contain an N-terminal AGENET domain (AGD) with four tandem AGD motifs and a C-terminal plant-specific DUF724 domain with unknown function. The AGENET domain-contain protein AGDP1 was reported to be required for transcriptional gene silencing and DNA methylation (Zhang et al., 2018). To determine whether transgene silencing in the *agdp3-1* and *agdp3-2* mutants is also associated with DNA methylation, we examined the DNA methylation levels of the *35S* promoter. Bisulfite sequencing showed that *35S* promoter was hypermethylated, especially in region A, in *agdp3-1* mutant (Supplemental Fig. 1a). To support the hypothesis that the transgene silencing was caused by increased DNA methylation, we treated the seedlings with the DNA methylation inhibitor 5-aza-2’-deoxycytidine. Not only the root phenotype, but the transgene silencing was also rescued in *agdp3-1* and *agdp3-2* mutants (Supplemental Fig. 1b and 1c).

To investigate whether AGDP3 also prevents endogenous loci from hypermethylation, we firstly used Chop PCR to determine DNA methylation level at the 3’ region of AT1G26400 and the promoter of AT4G18650, two genomic loci hypermethylated in *ros1* mutant plant (Qian et al., 2012). Both loci were also hypermethylated in *agdp3-1* plants (Supplemental Fig. 1d). The T-DNA insertion mutant allele *agdp3-3* in Col-0 background without *35S::SUC2* transgene also showed hypermethylation at both loci (Supplemental Fig. 1d). To understand the extent of *agdp3* mutation on genome-wide DNA methylation, we compared the methylomes of *agdp3-3* and wild-type Col-0 plants. In general, the overall DNA methylation patterns in genes and transposons were not altered by *agdp3* mutation.

However, we identified 1475 differentially methylated regions (DMRs) in the *agdp3-3* mutant plants compared to WT plants (Fig. 2a). The DMRs are distributed across each chromosome and about 32%, 39% and 27% of them are in the genic, TE and intergenic regions, respectively (Supplemental Fig. 2a and 2b). Among the 909 hyper-DMRs with significantly increased DNA methylation in *agdp3-3,* about 51% and 54% are also hypermethylated in *ros1* and *rdd* mutants, respectively. However, only 22% of the hyper-DMRs in *agdp3-3* overlap with those in *idm1-1* mutant (Fig. 2a and 2b). Together, these results indicate that AGDP3 regulates active DNA demethylation whole-genome wide in an IDM1-independent way.

**Figure. 2.**
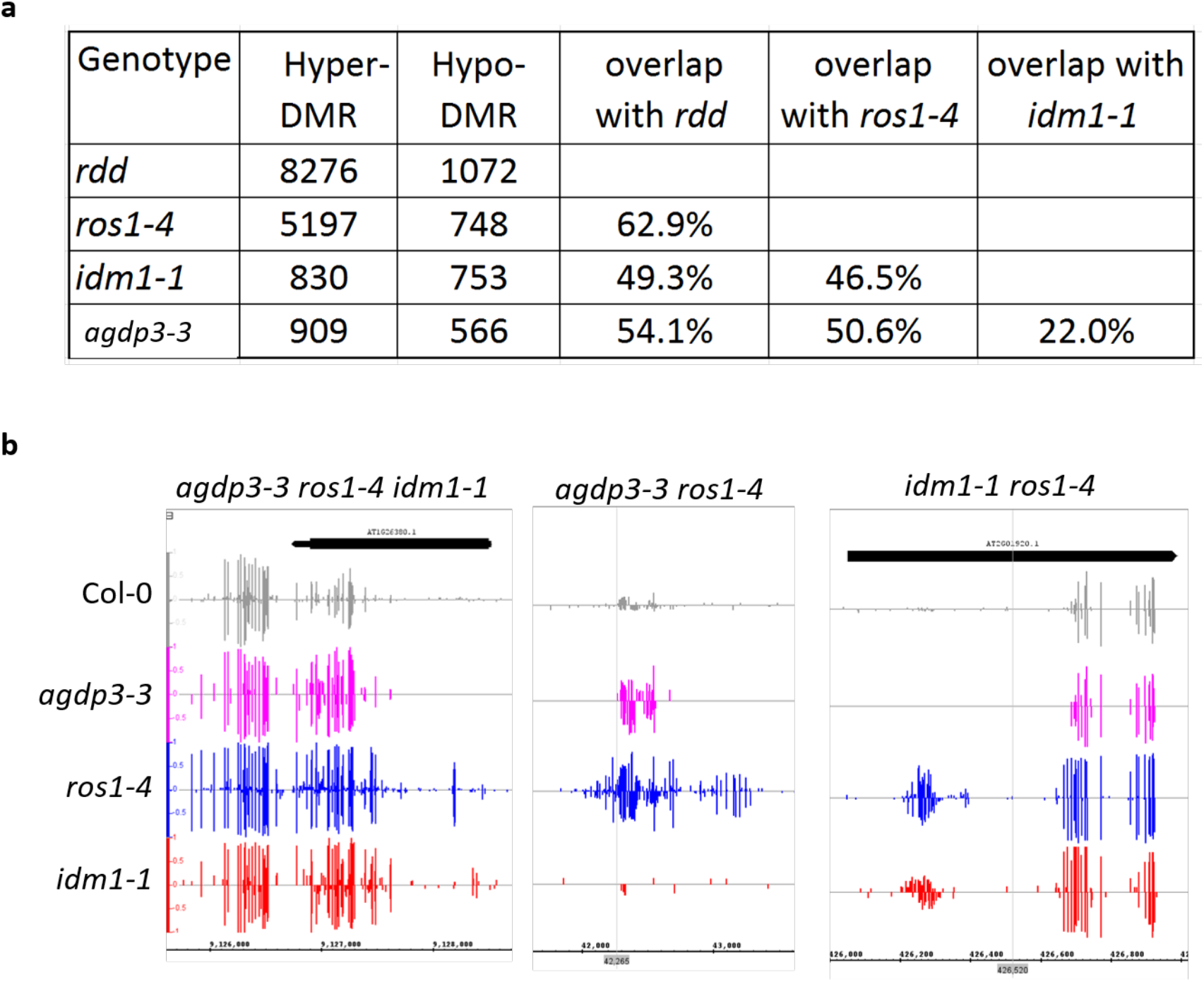
Analysis of DMRs identified in *agdp3-3* mutant. **a** The numbers of DMRs identified in *rdd*, *ros1-4, idm1-1* and *agdp3-3* and the overlap of hyper-DMRs between genotypes. **b** Examples of hyper-DMRs showing different pattern in *ros1-4, idm1-1* and *agdp3-3*.

### AGDP3 associates with genomic region of H3K9me2 mark

AGDP3 has an Agenet domain, suggesting that it may act as a histone reader to recognize some specific histone modification. The Agenet domains are plant-specific histone readers that recognize different histone modifications. To identify specific targets of AGDP3, we performed Cleavage Under Targets and Tagmentation (CUT&Tag) assay, using *agdp3-3* mutant transformed with *pAGDP3::AGDP3-3xFLAG,* which complemented the DNA hypermethylation phenotype. We identified 252 peaks of AGDP3 enrichment on chromatin, among which 218 (86.5%) overlapped with H3K9me2 peaks (Fig. 3). This result suggests that AGDP3 may recognize some specific loci with H3K9me2 mark and regulate DNA demethylation.

**Figure. 3.**
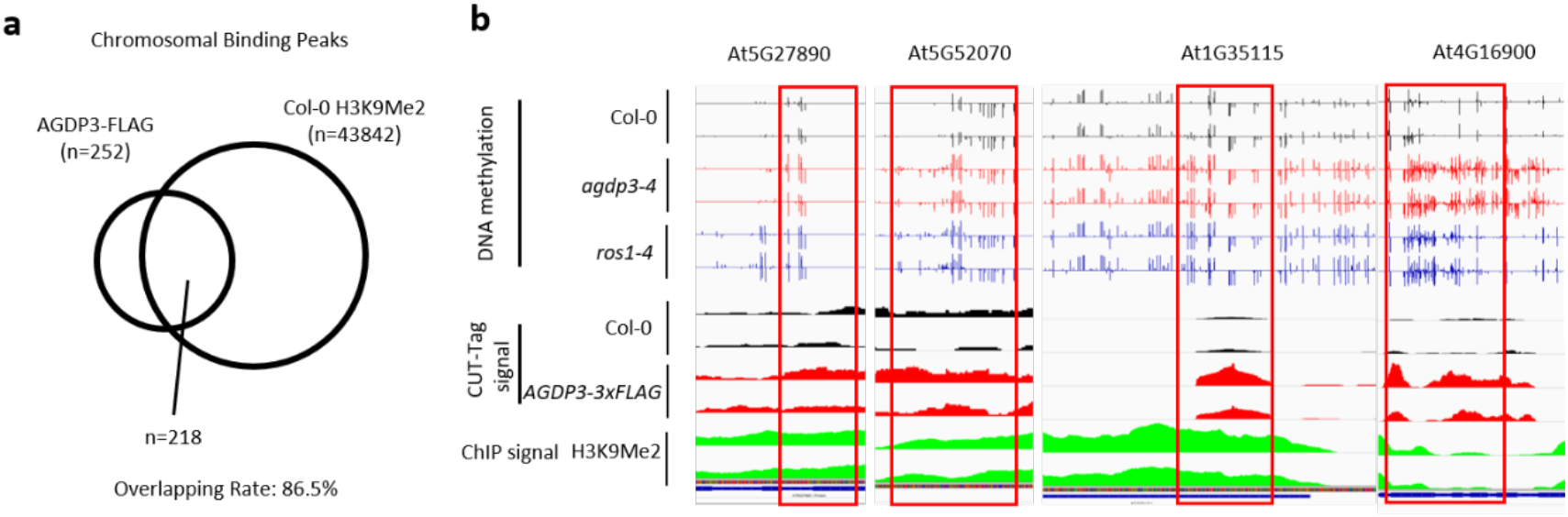
**a** Overlap of binding peaks between AGDP3 and H3K9Me2. **b** Examples of DNA methylation, AGDP3 enrichment and H3K9Me2 signal at several selected common hyper-DMRs in *ros1-4* and *agdp3-3.*

### AGDP3 AGD12 binds to H3K9me2 but AGD34 does not

The *Arabidopsis thaliana* AGDP3 (AtAGDP3) possesses four tandem AGDs (AGD1-4) at its N-terminus, which can be divided into two cassettes (AGD12 and AGD34) and a C-terminal Domain of Unknown Function 724 (DUF724) domain (Fig. 4a) (Marchler-Bauer et al., 2017). In plants, the tandem AGD cassette has been recently reported to function as a histone mark reader module to recognize the H3K9me2 mark (Chen, 2019; Harris and Jacobsen, 2019; Zhang et al., 2018a; Zhao et al., 2019). To explore the potential histone mark binding property of AGDP3, we expressed the AGD1-4 of AtAGDP3 and performed ITC-based *in vitro* binding assay (Supplemental Fig. 3a). The AGD1-4 of AtAGDP3 clearly shows a binding preference towards the H3K9me2 mark over other common histone marks (Supplemental Fig. 3a and 3b). The binding yields a binding ratio near 1 (Supplemental Fig. 3b), indicating that only one pair of tandem AGD cassettes is responsible for the H3K9me2 binding. To check which tandem AGDs recognize H3K9me2, we split the AGD1-4 to AGD12 and AGD34. However, both the isolated AGD12 and AGD34 of AtAGDP3 aggregated during purification and could not be used for further binding assay. Further, we chose its orthologs from other species for testing. We used the *Nicotiana tabacum* AGDP3 (NtAGDP3), which possesses the same domain architecture and shares 38% and 29% sequence identities with the AtAGDP3 AGD1-4 and DUF724 (Fig. 4a and Supplemental Fig. 4), respectively, for further experimentation. The NtAGDP3 AGD1-4 selectively binds to methylated H3K9 mark with a preference for H3K9me2, too (Fig. 4b and 4c). The AGD12 of NtAGDP3, but not AGD34, can specifically recognize the methylated H3K9 with a preference on the dimethylation, which is similar to the AtAGDP3 (Fig. 4d).

**Fig. 4.**
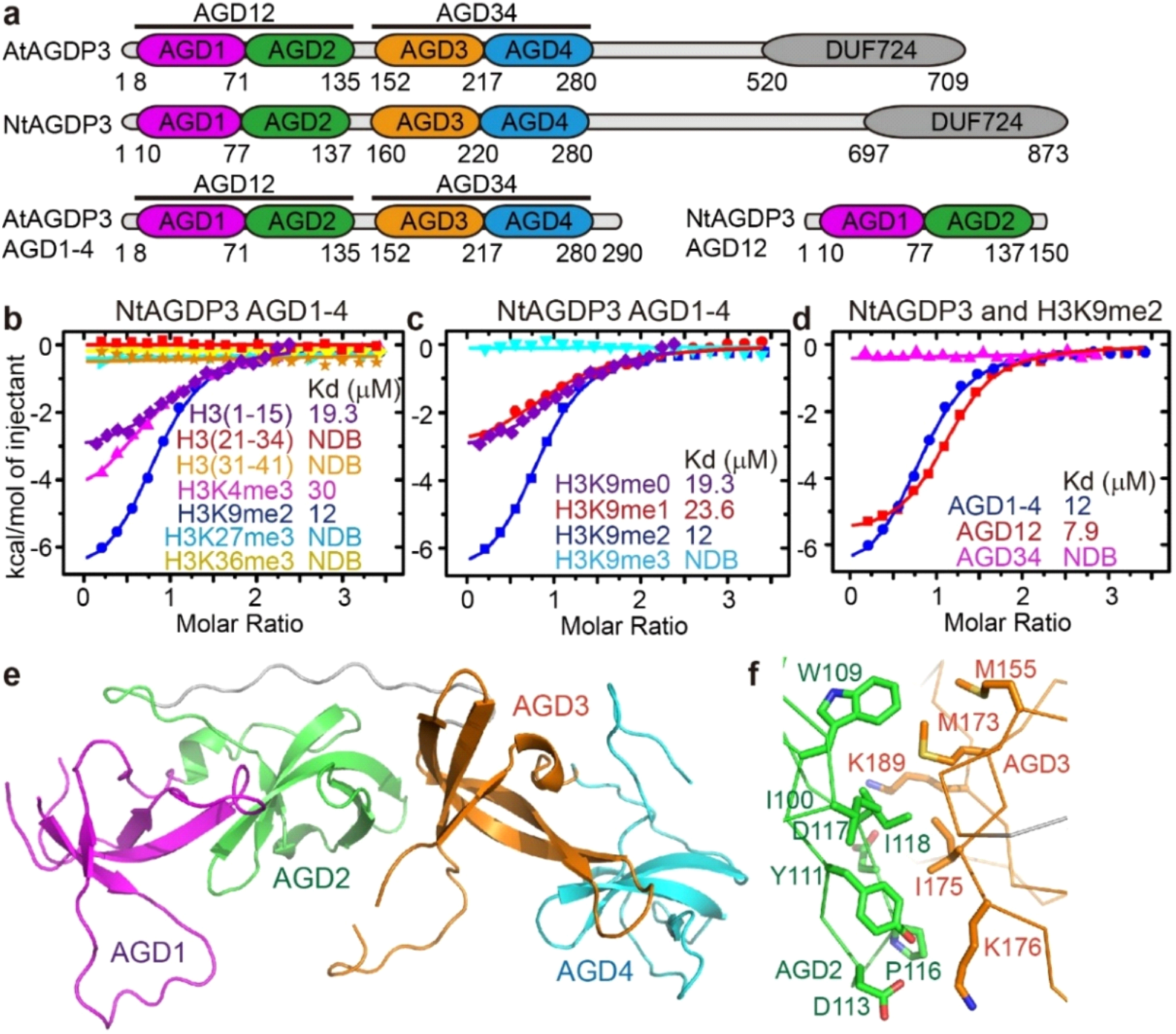
Overall structure of AGDP3 AGD1-4. **a** The schematic representation of the domain architecture of AtAGDP3 (upper panel), NtAGDP3 (middle panel), and the constructs used for structural studies (lower panel). **b-c** The ITC binding curves between NtAGDP3 AGD1-4 and various unmodified and methylated histone peptides (**b**) and different methylated status of H3K9 (**c**) reveal AGDP3 is an H3K9me2-specific reader. NDB, no detectable binding. **d** The ITC binding curves between different AGDP3 AGD cassettes and H3K9me2 reveal that the AGD12 cassette of AGDP3 is responsible for the H3K9me2 binding. **e** Overall structure of AtAGDP3 AGD1-4 with the four AGDs colored in magenta, green, orange, and cyan, respectively. **f** The interacting interface between AGD2 and AGD3 of AtAGDP3 with the interacting residues highlighted in stick models.

### Overall structure of AGD1-4

To investigate the molecular details of the recognition, we further performed structural studies. We determined the crystal structure of AtAGDP3 AGD1-4 and the structure was refined to 2.6 Å resolution (Fig. 4e and Table S1). The structure is composed of two tandem AGDs with AGD1 and AGD4 flanking on the two sides and AGD2 and AGD3 in the middle (Fig. 4E). Overall, both the AGD12 and AGD34 adopt classical tandem AGD folds resembling previously reported plant AGDP1 and human FMRP tandem AGDs (Myrick et al., 2015; Zhang et al., 2018a; Zhao et al., 2019b). The superimposition of AGDP3 AGD12 and AGD34 yields an RMSD of 2.0 Å for 127 aligned Cas, indicating quite similar overall structures. The interface between AGD2 and AGD3 is composed by a hydrophobic core and two pairs of salt bridge interactions on the sides (Fig. 4f).

### Structure basis for the recognition of H3K9me2 by AGD12 of AGDP3

To further investigate the molecular basis for the recognition of H3K9me2 by AGD12 of AGDP3, we tried to obtain the complex structure between AGDP3 and an H3K9me2 peptide. After extensive testing, we successfully got the crystal structure of NtAGDP3 AGD12 in complex with an H3(1-15)K9me2 peptide at 2.3 Å resolution (Fig. 5a and Table S1). The NtAGDP3 AGD12 resembles the AtAGDP3 AGD12 with an RMSD of 1.6 Å for 136 Cas upon superimposition (Fig. 5a). Similar to the AGDP1 AGD12-H3K9me2 complex, the peptide features a unique helical conformation that the H3K4 to H3A7 of the peptide forms a short a-helix (Fig. 5a). The peptide binds onto a negatively charged continuous surface between the two AGDs with the H3K4me0 and H3K9me2 side chains inserting into two surface pockets on the AGD1 and AGD2, respectively (Fig. 5a and 5b). The interaction between AGDP3 and H3K9me2 peptide focuses on three regions. At the N-terminus, the main chain carbonyl of AGD1 residue Ser55 forms two hydrogen bonds with the amino and amide groups of H3A1 and H3R2, respectively (Fig. 5c). The main chain carbonyl and side chain guanidine groups of H3R2 form hydrogen bonding and/or salt bridge interactions with Val57 of AGD1 and Asp99 of AGD2, respectively (Fig. 5c). The unmodified H3K4 forms salt bridge and hydrogen bonding interactions with Glu68 and Tyr53 of AGD1, respectively (Fig. 5d). Most importantly, the dimethyllysine of H3K9me2 is accommodated by an aromatic cage formed by Phe119, Trp102 and His97 (Fig. 5e), resembling other methyllysine readers (Patel, 2016). The mutations of the aromatic cage residues lead to the disruption of the binding between AGDP3 and H3K9me2, which is revealed by our ITC data (Fig. 5f). It is worth noting that all the NtAGDP3 residues involved in the interactions with H3K9me2 are strictly conserved in both sequence and structure with AtGADP3 (Fig. 6a) and the binding is similar to AGDP1 (Fig. 6b) (Zhang et al., 2018a; Zhao et al., 2019). However, these H3K9me2 interacting residues are not conserved in the AGD34 of AtAGDP3 structurally (Fig. 6c), consistent with our biochemical data that AGD12 but not AGD34 of AGDP3 recognizes H3K9me2 mark. Meanwhile, the AGD34 may have functions other than the H3K9me2 binding, which remains to be elucidated in the future.

**Fig. 5.**
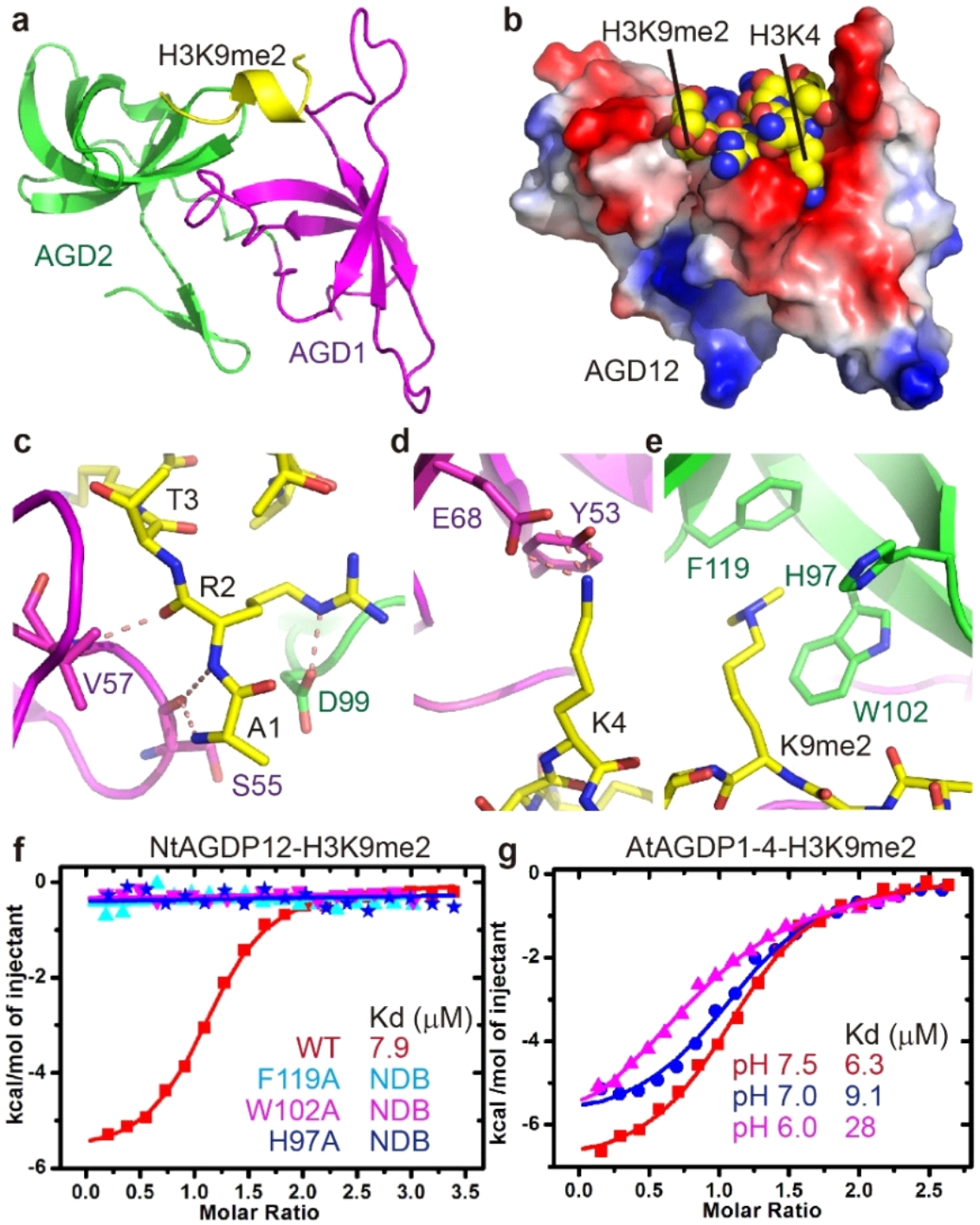
Structure of NtAGDP3 AGD12 in complex with an H3K9me2 peptide. **a** A ribbon diagram of the overall structure of NtAGDP3 AGD12-H3K9me2 complex with the AGD1, AGD2, and the peptide colored magenta, green, and yellow, respectively. The H3K9me2 peptide adopts a helical conformation. **b** An electrostatic surface view of the NtAGDP3 AGD12 with the H3K9me2 peptide in space filling model. The unmodified H3K4 and dimethylated H3K9 insert their side chains into two adjacent surface pockets of AGD12. **c-e** The detailed interaction between AGD12 and the peptide residues H3A1 and H3R2 (**c**), H3K4 (**d**), and H3K9me2 (**e**). The interacting residues and hydrogen bonds are highlighted in stick and dashed red lines, respectively. **f** The ITC binding curves between different AGD12 mutations that disrupt the H3K9me2 binding aromatic cage and H3K9me2 peptide revealing the aromatic cage is essential for the peptide binding. **g** The ITC binding curves between AtAGDP3 AGD1-4 and H3K9me2 peptide in different pH indicating AGDP3 is a pH-dependent H3K9me2 binding module. NDB, no detectable binding.

### AGDP3 is a pH sensitive H3K9me2 reader

Sine the pKa of the protonated imidazole group of the histidine side chain is around 6.0-7.5 in folded proteins and the plant cell nucleus has a pH of 7.2 ± 0.2 (Shen et al., 2013), the existence of a histone residue in the methyllysine binding aromatic cage raises the possibility that the binding of H3K9me2 by AGDP3 is pH dependent. As the AGD12 of NtAGDP3 precipitates heavily in low pH, we used AtAGDP3 AGD1-4 for testing. Consequently, the protonated histidine containing aromatic cage of AtAGDP3 showed about 5-fold decreased binding towards H3K9me2 in pH 6.0 than in pH 7.5 (Fig. 5g). In contrast, the H90Y mutant of AtAGDP3, which corresponds to H97Y of NtAGDP3, loses the pH sensitivity (Fig. 6d), confirming that the histidine residue of the aromatic cage is responsible for the pH-dependent binding of H3K9me2 by AGDP3. This type of pH dependent binding of histone mark is similar to the recognition of H3K4me3 by the *Drosophila* Protein Partner of Sans-fille (PPS) and the recognition of H3K122suc by Glioma-Amplified Sequence-41 (GAS41) (Tencer et al., 2017; Wang et al., 2018), indicating this might be a common regulatory mechanism for histone readers.

**Fig. 6.**
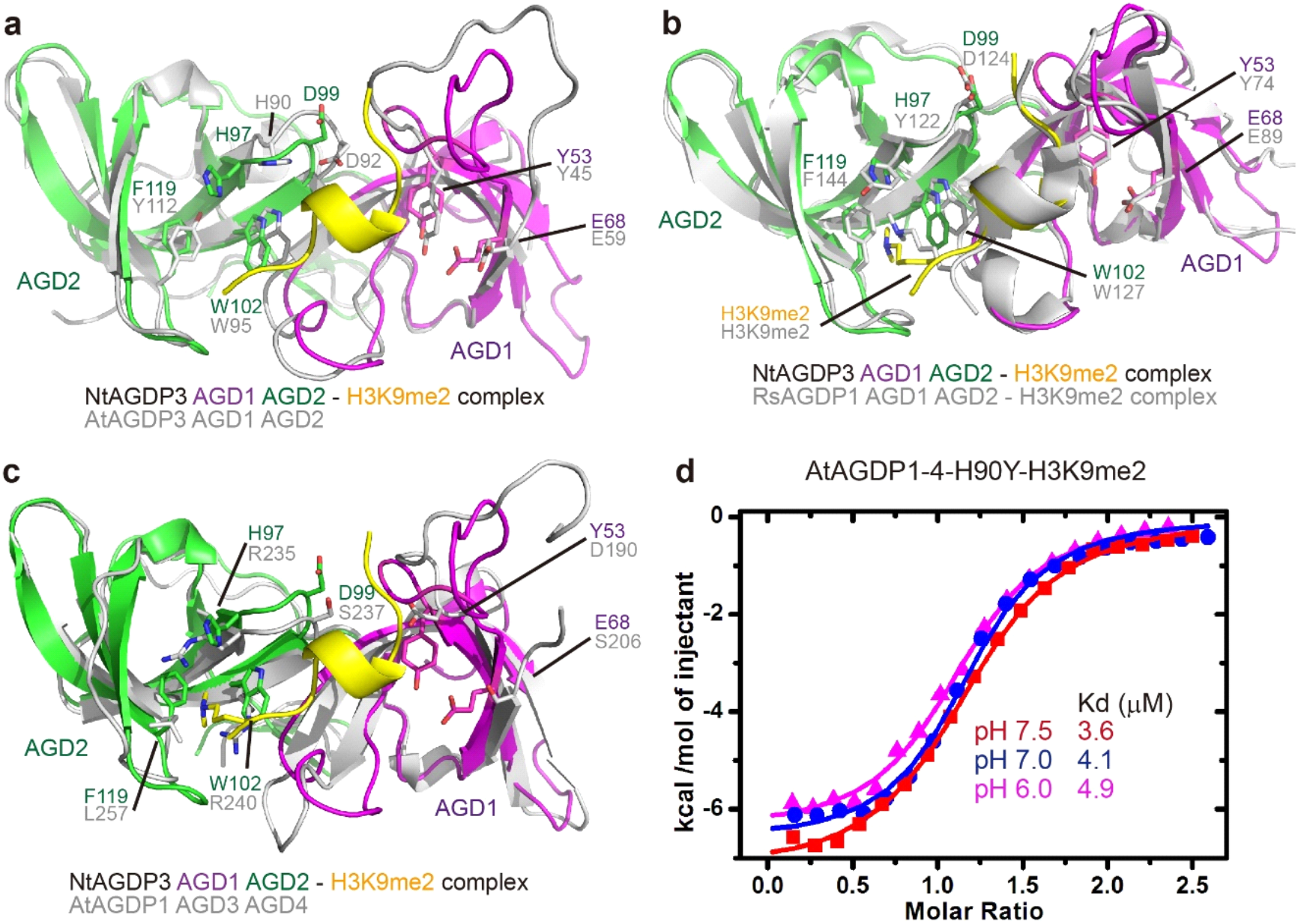
The structural comparison of different tandem AGDs and AGDP3 is a pH dependent H3K9me2 reader. **a** The superimposition of NtAGDP3 AGD12-H3K9me2 with the free form AtAGDp3 AGD12 shows almost identical conformation and key residue positions, suggesting that AtAGDP3 AGD12 recognizes H3K9me2 in a similar way as NtAGDP3 AGD12. **b** The superimposition of NtAGDP3 AGD12-H3K9me2 complex and RsAGDP1 AGD12-H3K9me2 complex shows similar conformation and key residue positions, indicating similar H3K9me2 recognition mechanism. The major difference is that the aromatic residues Tyr122 in RsAGDP1 is replaced by His97 in NtAGDP3. **c** The superimposition of NtAGDP3 AGD12 and AtAGDP3 AGD34 shows that the aromatic cage residues of AGD12 is not conserved in AGD34, suggesting AGD34 is not a methyl lysine reader and may possess other functions. **d** The ITC measurement of AtAGDP3 AGD1-4 H90Y mutant with H3K9me2 in different pH shows similar binding affinity, suggesting that the histidine residues is responsible for the pH dependence.

## Discussion

Active DNA demethylation is a dynamic process that is manipulated by many different regulatory factors and enzymes that work coordinately. Even though ROS1 has been extensively studied in enzymology, the mechanisms of recruitment to its target sites are still poorly understood in plants. In this study, we isolated the AGENET domain containing protein AGDP3 as an anti-silencing factor from a forward genetic screen, and find that ADGP3 functions as a cellular DNA demethylation regulator that inhibits DNA hypermethylation at multiple genomic regions. Our results indicate that AGDP3 prevents genome-wide DNA methylation and protects endogenous genes from silencing, showing important roles in epigenetic regulation and active DNA demethylation. Epigenetic modification enzymes are usually associated with DNA- or chromatin-binding proteins or domains for specific targeting (DesJarlais and Tummino, 2016). The current study suggests that ADGP3 may form a protein complex with ROS1 to function in active DNA demethylation. So far, two different active DNA methylation pathways have been identified in Arabidopsis: H2A.Z-dependent recruitment of ROS1 to specific genomic regions binding by IDM1 and MBD7 in the IDM complex (Nie et al., 2019; Qian et al., 2012) and RWD40-dependent recruitment of ROS1 to specific target loci binding by RMB1 in RWD40 complex (Liu et al., 2020). However, it should be noted that even additive DMRs contributed by these two pathways are much fewer than the hyper-DMRs induced by dysfunction of ROS1, which indicates the complexity of active DNA methylation in plants. The ADPG3 identified in this study contributes to a subset of genomic methylated regions targeted by ROS1, which is different from the IDM complex- or RWD40 complex-mediated pathway. Hence, our study enriches the mechanisms of active DNA demethylation in plants.

Recently, the crystal structures of tandem AGDs of AGDP1 in complex with H3K9me2 were reported (Zhang et al., 2018a; Zhao et al., 2019). The superimposition of AGDP1 AGD12-H3K9me2 complex and AGDP3 AGD12-H3K9me2 complex yielded an RMSD of 1.2 Å for 136 Cas, revealing almost identical overall structures (Fig. 6b). Most of the key residues involved in peptide recognition are strictly conserved, and the most obvious difference is the aromatic cage residue His97 in NtAGDP3 which is equivalent to the Tyr122 residue in RsAGDP1 (Fig. 6b). The existence of a histidine residue in the aromatic cage enables AGDP3 pH dependent binding of H3K9me2 (Fig. 5g). As AGDP1 is considered as a plant functional analog of animal HP1 protein and is tightly associated with heterochromatic H3K9me2 and DNA methylation (Zhang et al., 2018a; Zhao et al., 2019), it is convictive for AGDP1 to possess constitutive binding towards the gene silencing mark H3K9me2, no matter how pH changes. In contrast to DNA methylation, DNA demethylation occurs more dynamically as an on-or-off switch and requires the changing of the micro-epigenetic environment from silencing to activation. Especially, the AGDP3 associates with H3K9me2 but not the ROS1 associated H3K18ac and H3K27me3 marks, indicating that AGDP3 targeting to chromatin is more dynamically regulated.

It was reported that global histone acetylation, a signal of gene activation, is associated with high intracellular pH (pH 7.4), while histone deacetylation is with low intracellular pH (pH 6.5) in animal cells (McBrian et al., 2013). Considering the common relationship between histone modification and metabolism, we think the plant cell may possess a similar phenomenon. Therefore, we can build a plausible molecular model for the action of AGDP3 in ROS1-mediated anti-silencing (Fig. 7). Once the chromatin loci are marked by the gene activation signal of histone acetylation, probably including the H3K18ac, the histone acetylation-associated higher pH enables AGDP3 to specifically bind to H3K9me2 marked chromatin loci and to further recruit ROS1 to mediate DNA demethylation for encountering gene silencing (Fig. 7). In the gene silencing process, the dropping of histone deacetylation-associated pH may disrupt the binding between AGDP3 and the chromatin, which may further prevent the ROS1 anti-silencing machinery to approach (Fig. 7), thereby keeping the DNA methylation mark and maintaining gene silencing chromatic states. Although there is no evidence that the histone acetylation can directly recruit ROS1 anti-silencing complex or not, it can increase the pH of the microenvironment of chromatin, which further enables H3K9me2 binding by AGDP3. Then, AGDP3 associated ROS1 may subsequently specifically demethylate DNA, removing the gene silencing mark. Overall, we speculate that AGDP3 may function as an indirect histone acetylation sensor via direct sensing of the pH to further connect ROS1 anti-silencing machinery with its substrate, H3K9me2/DNA methylation marked silencing chromatin loci. Our work sheds light on the potential internal crosstalk among histone acetylation, H3K9me2 and DNA methylation.

**Fig. 7.**
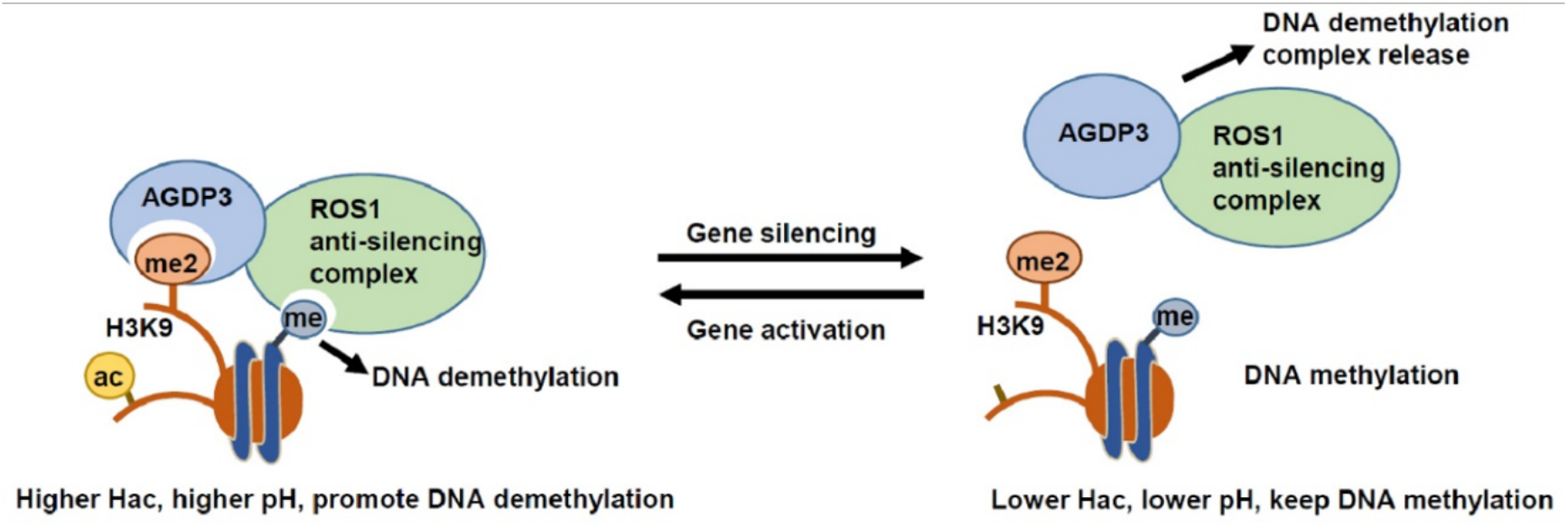
A schematic model for the molecular function of AGDP3. In gene activation loci, the active mark histone acetylation-associated high pH may enable the binding of H3K9me2 by AGDP3, which further recruits ROS1 anti-silencing complex through an unknown factor to remove DNA methylation (left panel). In contrast, in gene silencing loci, the histone deacetylation associated low pH may be sensed by AGDP3 and further release the H3K9me2 binding, allowing the ROS1 anti-silencing complex to be released and keep DNA methylation (right panel).

## Methods

### Plant materials, mutant screening and map-based cloning

Wild-type (WT) in this study were described previously (Lei et al., 2011). An EMS-mutagenized pool of plants was generated and screened for mutants with a long-root phenotype (Wang et al., 2013). M2 seedlings were grown vertically on ½ MS plates with 2% sucrose and 1% agar and mutants with long-root phenotype among 7-day-old seedlings were screened. Genetic mapping and gene cloning was performed as described previously (Lei et al., 2014).

### Chop-PCR

Chop-PCR assays were carried out according to Lei et al (2014).

### Whole-genome bisulfite sequencing and data analysis

Genomic DNA was extracted from 14-day-old seedlings using Dneasy Plant Maxi Kit (Qiagen). Bisulfite conversion, library construction, and deep sequencing were performed by the Novogene Co., Ltd. in Beijing, China. DMRs were identified according to Qian et al. (2012) with some modifications.

### Cleavage Under Targets and Tagmentation (CUT&TAG) Assay

Experiments were carried out using Hyperactive In-Situ ChIP Library Prep Kit for Illumina (pG-Tn5) (Vazyme Biotech Co.,Ltd). Briefly, ~1g flower tissue of pAGDP3-AGDP3-3xFLAG/agdp3-3 transgenic plants were collected and directly flash frozen in liquid nitrogen and grinded into fine powder after harvest. Cell nucleus were isolated by filtering through 4 layers of miracloth. Following steps were performed as described in the manufacturer’s mannual. The WGS library was sent to the Novogene Co., Ltd. in Beijing, China for Illumina sequencing.

### Protein expression and purification

The gene encoding NtAGDP3 AGD12 (1-150) was cloned into a pSumo vector to fuse an N-terminal hexahistidine plus yeast Sumo tag to the target proteins. The plasmid was transformed into the *E. coli* strain BL21 (DE3) RIL and cultured in LB medium. The protein expression was induced by adding IPTG to a final concentration of 0.15 mM when the OD600 of cell culture reached 0.6. The recombinant expressed protein was purified using HisTrap column (GE Healthcare). The His-Sumo tag was digested by ulp1 protease and removed by a second HisTrap column. The protein was further purified using a Heparin column (GE Healthcare) and a Superdex G200 column (GE Healthcare). The AtAGDP3 AGD1-4 (1-290), AGD12 (1-143), AGD34 (153-290), NtAGDP3 AGD1-4 (1-296) and AGD34 (150-296) were cloned, expressed and purified using the same protocol. The site-directed mutagenesis was conducted using a PCR-based method. All the mutant proteins were expressed and purified using the same protocol as the wild type protein. All the peptides were purchased from GL Biochem Ltd (Shanghai).

### Crystallization and structure determination

The crystallization was carried out using vapor diffusion sitting drop method at 20 °C. To facilitate crystallization, a surface entropy reduction method was applied to AtAGDP3 AGD1-4 to generate suitable mutations for crystallization (Goldschmidt et al., 2007). The AtAGDP3 AGD1-4 K261A/K262A/E263A mutant was crystallized in a condition of 0.1 M lithium sulfate, 20% PEG1000 and 0.1M sodium citrate, pH 5.5. For protein-peptide complex formation, the purified NtAGDP3 AGD12 was mixed with an H3(1-15)K9me2 peptide with a molar ratio of 1:3 for 2 hours at 4 °C. The complex crystal was obtained in a condition of 25% PEG 1500, 0.1 M sodium chloride and 0.1 M bis-Tris propane, pH 9.0. To resolve the phase by heavy atoms, an AtAGDP3 AGD1-4 crystal was soaked in the reservoir solution supplemented with 5 mM ethylmercurithiosalicylic acid at 20 °C for 2 h. All the crystals were cryo-protected in the reservoir solution supplemented with 20% glycerol and then flash cooled into liquid nitrogen. All the diffraction data were collected at beamline BL19U1 of the National Center for Protein Sciences Shanghai (NCPSS) at the Shanghai Synchrotron Radiation Facility (SSRF) and further processed using the program HKL3000 (Otwinowski and Minor, 1997). A summary of the statistics of the data collection is listed in **Table S1**.

The structure of AtAGDP3 AGD1-4 was determined using single-wavelength anomalous diffraction (SAD) method with the mercury derivate data using the program Phenix (Adams et al., 2010). The structure refinement and model building were applied using the programs Phenix and Coot, respectively (Adams et al., 2010; Emsley et al., 2010). The structure of NtAGDP3 AGD12 in complex with H3K9me2 peptide was determined using molecular replacement method using the program Phenix with the structure of RsAGDP1 AGD12-H3K9me2 complex as search model (Adams et al., 2010; Zhang et al., 2018a). The NtAGDP3 AGD12-H3K9me2 complex data were anisotropic and were truncated by the Diffraction Anisotropy Server (https://services.mbi.ucla.edu/anisoscale/) (Strong et al., 2006). The structure refinement and model building were performed using the same protocol as AtAGDP3 AGD1-4. A summary of the structure refinement statistics is listed in **Table S1**. The molecular graphics were generated using the program Pymol (Schrödinger, LLC). The sequence alignments were performed using the program T-coffee and illustrated using the ESPript server (Di Tommaso et al., 2011; Robert and Gouet, 2014).

### ITC

The *in vitro* binding assays were performed using a Microcal PEAQ ITC instrument (Malvern). The sample proteins were dialyzed into a titration buffer of 100 mM NaCl and 20 mH HEPES, pH 7.5. For the pH dependence test, the low pH buffer of 100 mM NaCl, 20 mM sodium/potassium phosphate, pH 7.0 or pH 8.2 were used. The peptides were dissolved into the same buffer. The titration experiments were performed at 20 °C and the data were fit using the program Origin 7.0.

## Data availability

X-ray structures have been deposited in the RCSB Protein Data Bank with the accession codes: XXXX for the AtAGDP3 AGD1-4 and XXXX for NtAGDP3 AGD12 in complex with H3K9me2 peptide.

## Author contributions

X.Z., J.L., X.D., R.L. and Z.Y. performed the structural and biochemical experiments. M.W., W.N., Y.X., V.S., H.L., Y.Z., and Y.Y., performed genetic experiments. K.T., L.P., and Q.Z. did the bioinformatic analysis. J.Z., J.D. and M.L. conceived the study, designed the experiments and wrote the paper.

## ACKNOWLEDGMENTS

We thank the staffs at beamline BL19U1 of the National Center for Protein Sciences Shanghai (NCPSS) at the Shanghai Synchrotron Radiation Facility (SSRF) for the X-ray data collection. This work was supported by the Chinese Academy of Sciences and National Natural Science Foundation of China (31970580) to M.L., and by National Key R&D Program (2016YFA0503200), Shenzhen Science and Technology Program (JCYJ20200109110403829 and KQTD20190929173906742) and Key Laboratory of Molecular Design for Plant Cell Factory of Guangdong Higher Education Institutes (2019KSYS006) to J.D.

