## supplemental data for "A H3K9me2-Binding Protein AGDP3 Limits DNA Methylation and Transcriptional Gene Silencing in Arabidopsis"

### Supplementary Figures

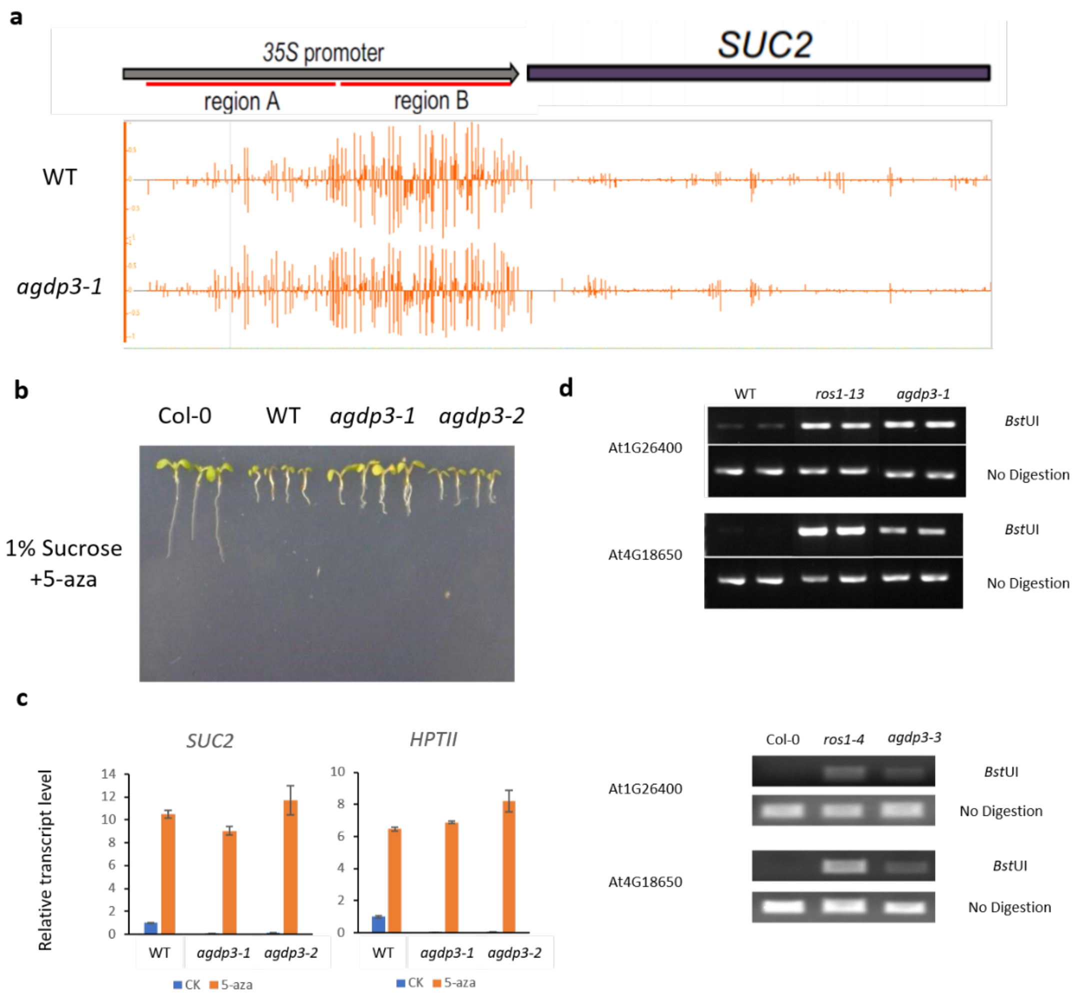


**Supplemental Figure. 1. a** DNA methylation status at the promotor of *35S::SUC2* transgene in WT and *agdp3-1*. **b** Short root phenotype of *agdp3-1* and *agdp3-2* could be restored by a DNA methylation inhibitor, 5-Aza-2'-deoxycytidine treatment. **c** 5-Aza-2'-deoxycytidine treatment restored the transcripts level of transgenes in *agdp3-1* and *agdp3-2*. **d** CHOP-PCR verification of DNA hyper-methylation in *agdp3-1*, *agdp3-2* and *agdp3-3*.

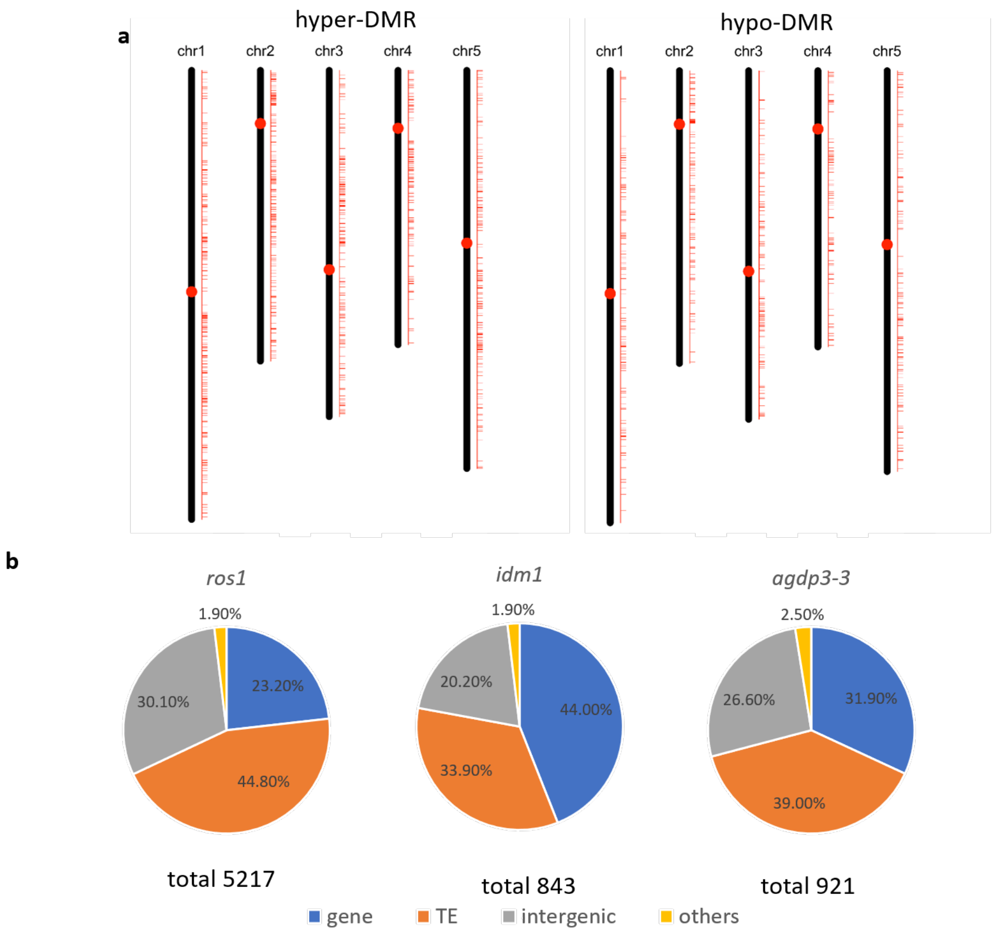


**Supplemental Figure. 2. a** Chromosomal distribution of DMRs in *agdp3-3*. **b** Composition of *ros1-4*, *idm1-1* and *agdp3-3* hyper-DMRs.

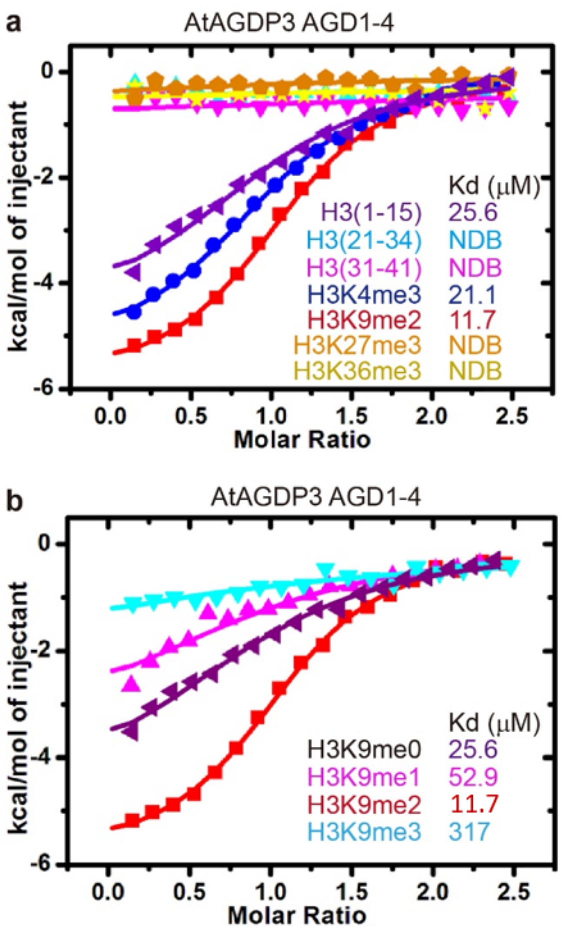


**Supplementary Fig. 3.** AtAGDP3 specifically recognizes H3K9me2 mark. **a-b** The ITC binding curves between AtAGDP3 AGD1-4 and various unmodified and methylated histone peptides (**a**) and different methylated status of H3K9 peptides (**b**) revealing AtAGDP3 can specifically recognize the H3K9me2 mark.

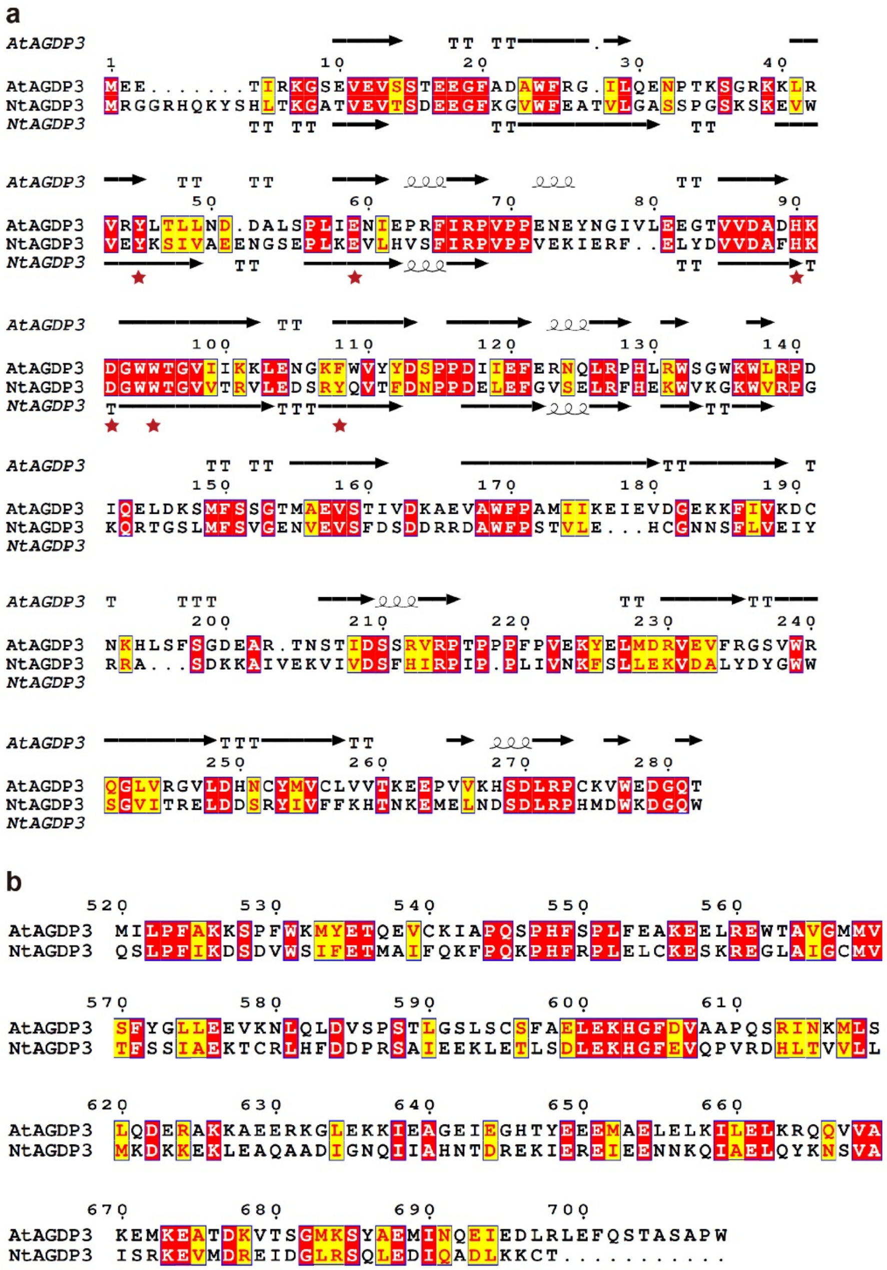


**Supplementary Fig. 4.** Sequence alignment of AtAGDP3 and NtAGDP3. **a** Structure-based sequence alignment of the AGD1-4 of AtAGDP3 and NtAGDP3. The secondary structures of AtAGDP3 and NtAGDP3 are highlighted on the top and bottom of the alignment. The residues involving peptide recognition are marked by red stars and are conserved. **b** Sequence alignment of the DUF724 domain of AtAGDP3 and NtAGDP3 showing that the two domains are conserved.

**Table S1. Data collection and refinement statistics**

| | AtAGDP3 AGD1-4 Hg | AtAGDP3 AGD1-4 Native | NtAGDP3 AGD12 with H3K4me3 |
|---|---|---|---|
| **Data collection** | | | |
| Beamline | SSRF-BL19U1 | SSRF-BL19U1 | SSRF-BL19U1 |
| PDB code | | XXXX | XXXX |
| Space group | $I2_12_12_1$ | $I2_12_12_1$ | $P4_12_12$ |
| Wavelength (Å) | 0.9789 | 0.9789 | 0.9789 |
| Cell dimensions | | | |
| *a*, *b*, *c* (Å) | 78.3, 129.8, 168.3 | 78.5, 129.2, 167.0 | 70.7, 70.7, 218.0 |
| Resolution (Å) | 50.0-3.9 (4.04-3.90) [a] | 50.0-2.6 (2.69-2.60) | 50.0-2.3 (2.38-2.30) |
| $R_{merge}$ | 0.126 (0.951) | 0.039 (0.705) | 0.090 (0.979) |
| $I / \sigma I$ | 36.0 (10.2) | 30.4 (1.6) | 32.1 (2.7) |
| Completeness (%) | 99.8 (100.0) | 99.2 (99.7) | 100.0 (100.0) |
| Redundancy | 12.8 (13.1) | 4.4 (4.5) | 24.6 (21.0) |
| **Refinement** | | | |
| $R_{work}$ / $R_{free}$ | | 0.231 / 0.260 | 0.232 / 0.268 |
| No. reflections | | 26,331 | 19,193 |
| No. atoms | | 4,425 | 3,691 |
| Protein / Peptide | | 4,419 / - | 3,383 / 238 |
| Water | | 6 | 70 |
| *B*-factors ($Å^2$) | | 94.6 | 58.2 |
| Protein / Peptide | | 94.6 / - | 57.0 / 80.1 |
| Water | | 68.9 | 43.0 |
| R.m.s. deviations | | | |
| Bond lengths (Å) | | 0.010 | 0.004 |
| Bond angles (°) | | 1.231 | 0.966 |

[a] Highest-resolution shell is shown in parentheses.
